# Rethinking inheritance, yet again: inheritomes, contextomes and dynamic phenotypes

**DOI:** 10.1101/013367

**Authors:** N. G. Prasad, Sutirth Dey, Amitabh Joshi, T. N. C. Vidya

## Abstract

In recent years, there have been many calls for an extended evolutionary synthesis, based in part upon growing evidence for non-genetic mechanisms of inheritance, i.e., similarities in phenotype between parents and offspring that are not due to shared genes. While there has been an impressive marshalling of evidence for diverse forms of non-genetic inheritance (epigenetic, ecological, behavioural, symbolic), there have been relatively few studies trying to integrate the different forms of inheritance into a common conceptual structure, a development that would be important to formalizing elements of the extended evolutionary synthesis. Here, we propose a framework for an extended view of inheritance and introduce some conceptual distinctions that we believe are important to this issue. In this framework, the phenotype is conceived of as a dynamic entity, its state at any point in time resulting from intertwined effects of previous phenotypic state, and of hereditary materials (DNA and otherwise) and environment. We contrast our framework with the standard gene-based view of inheritance, and also discuss our framework in the specific context of recent attempts to accommodate non-genetic inheritance within the framework of classical quantitative genetics and the Price equation. In particular, we believe that the extended view of inheritance and effects on the phenotype developed here is particularly well-suited to individual-based simulation studies of evolutionary dynamics. The results of such simulations, in turn, could be useful for assessing how well extended models based on quantitative genetics or the Price equation perform at capturing complex evolutionary dynamics.

## Introduction

In recent decades, increasing attention within evolutionary biology is being focused on the issue of non-genetic (i.e. not based on gene sequence variation) inheritance (reviewed at length in Jablonka and Lamb 2005; Pigliucci and Müller 2010). There is increasing evidence for the inheritance of environmentally induced epigenetic states from parents (reviewed in Bonduriansky and Day 2009; Jablonka and Raz 2009), as well as for the passing on of culturally acquired (i.e. learnt) behaviours to offspring and, indeed, other peers, referred to as cultural inheritance or transmission (Cavalli-Sforza and Feldman 1981; Richerson and Boyd 2005; Jablonka and Lamb 2005; El Mouden et al. 2014). The literature in these areas is growing but is also often confusing with regard to terminology and underlying concepts as a result of arising from diverse subdisciplinary backgrounds of inquiry. In this brief essay, we delineate some concepts and conceptual distinctions, and attendant terminology, that we believe may help ameliorate some of the confusion in attempts to bring together various forms of inheritance into a coherent and expanded view of evolutionary dynamics. Much of what we are drawing upon has been said before: what we hope to accomplish is to (a) highlight some specific aspects of present attempts to extend the evolutionary synthesis with regard to non-genetic inheritance where we believe that conceptual clarity is still lacking, and (b) make some suggestions as to how that conceptual clarity can be attained. Given the numerous recent reviews of different aspects on non-genetic inheritance and its evolutionary implications, the list of references is not comprehensive and is somewhat biased towards broad reviews or books wherein leads to most of the primary literature can be found.

Following the rediscovery of Mendel’s laws, the study of heredity gradually narrowed to an almost exclusive focus on genic inheritance, especially after Johannsen’s (1911) delineation of the concepts of genotype and phenotype, mirroring the hard dichotomy of germline and soma laid down by Weissmann (1904), that excluded any possibility of the inheritance of acquired characteristics (discussed in Schwarz 2008). Subsequently, this narrower, Mendelian, version of heredity was combined with an emphasis on natural selection as the principal mechanism of adaptive evolution, largely through the work of Fisher (1930), Haldane (1932) and Wright (1932), becoming a major foundation of the Modern Synthesis (Mayr 1992). The importance of genes as explanatory factors in biology got further strengthened following the advent of molecular genetic understanding of DNA structure, replication, expression and regulation. Thus, as heredity was central to the mechanism of natural selection, the narrowing of heredity to gene-based inheritance naturally led to the gene-centric bias of much modern evolutionary thinking, at least insofar as it pertained to adaptive evolution within populations (i.e. microevolutionary change) (discussed in Gould 2002; Amundson 2005; Bonduriansky 2012).

While the gene-centric version of microevolutionary theory has been extensively criticized for being limited in scope, especially in the light of ever-increasing evidence for non-genetic inheritance (e.g. Richerson and Boyd 1985; Odling-Smee et al. 2003; Dey and Joshi 2004; Jablonka and Lamb 2005; Pigliucci and Müller 2010), certain aspects of this theory, we believe, have not received as much attention from critics as they should have. First, in addition to the framework of population genetics in the strict sense, there is also the essentially phenotypic framework of quantitative genetics, arising from Fisher’s (1918) seminal paper on the correlations between relatives for polygenic traits following Mendelian inheritance. Interestingly, several critics of the Modern Synthesis ignore the quantitative genetics tradition altogether (e.g. Amundson 2005; Jablonka and Lamb 2005; Laland et al. 2014), although it is considerably more relevant than population genetics is to understanding adaptive evolution. While quantitative genetics did attempt to root its statistical analysis of trait correlations and evolution in Mendelian genetics, its most important insights – the notion of breeding value for fitness, and its variance, as determinants of adaptive evolutionary responses to fitness differences among individuals – are essentially not dependent on any underlying mechanistic model of inheritance. Indeed, the statistical approach of quantitative genetics, which effectively black-boxed the details of how genotypes affect phenotypes, reached its later, even more generalized, fruition in the Price (1970, 1972) equation that makes no assumption about the pattern of inheritance at all, subsuming it into a correlation of phenotypic values between individuals and immediate descendants. The results from this statistical tradition suggest that what ultimately matters in determining the response to a given selective scenario is not so much the genes that pass from parent and offspring but rather their statistical effects on offspring phenotype. This important distinction stems from the fact that heredity is ultimately about phenotypic similarity between parents and offspring, even though one major underlying reason for such similarity is the material genes that pass from parent to offspring, albeit subject to the vagaries of mutation, recombination, sex and chance. Thus, what an individual inherits, in terms of a genetic endowment leading to the propensity towards certain phenotypic values, is usually different from what the same individual transmits to its offspring in terms of expected effects on the offsrping phenotype. This principle is reflected in the clear distinction between the genotypic value (G) and the breeding value (A) of a trait in quantitative genetics (Falconer and Mackay 1996). This distinction has implications for the development of an extended theory of evolutionary change incorporating varied non-genetic forms of inheritance. There are now some studies that use the framework of quantitative genetics or the Price equation to incorporate non-genetic inheritance into models of evolutionary change (Bonduriansky and Day 2009; Helanterä and Uller 2010; Danchin et al. 2011; Santure and Spencer 2011; El Mouden et al. 2013), and we will discuss them in subsequent sections.

A second aspect of the gene-centric view of the evolutionary process that we believe has not received much critical attention is the essentially static conception of genotype and phenotype, at least within an individual’s lifetime, as also pointed out by Lewontin (2002) and Dey and Joshi (2004). Even the notion of the genotype-phenotype (G-P) map (Lewontin 1974: Fig. 1, pg. 14), which was later used to argue against the black-boxing of the details of development in Mendelian genetics (Alberch 1991), is an attempt to define a relationship (a map) linking genotypic space and phenotypic space. Despite the complexity of this relationship, it is nevertheless conceived of as a relationship between two spaces that are static within an individual’s lifetime, though not over evolutionary timescales.

**Figure 1.**
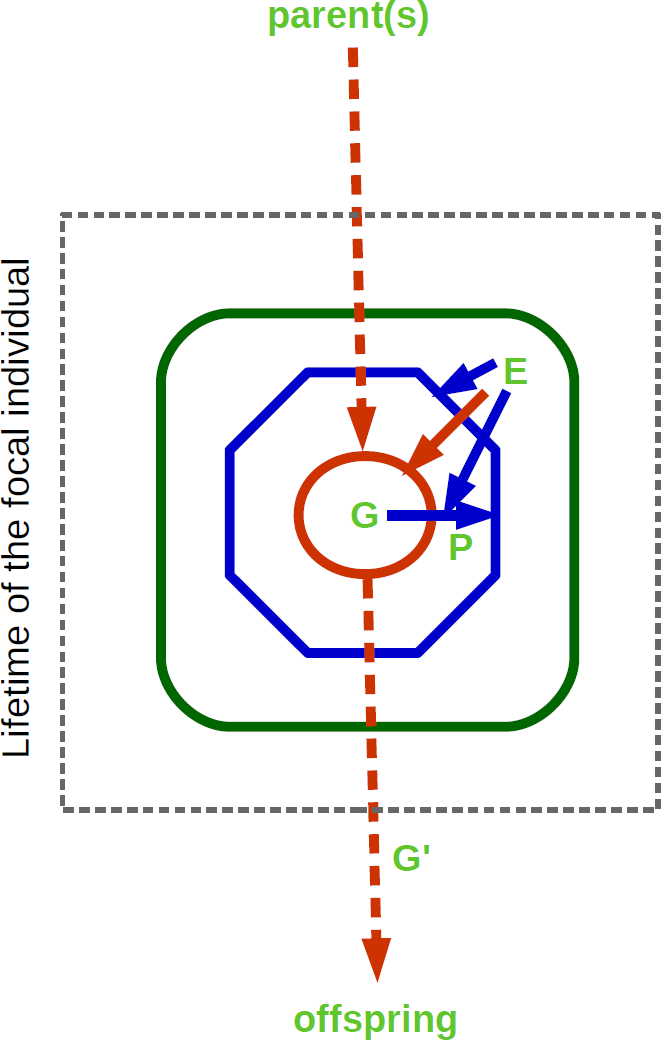
Schematic representation of the relationships between phenotype, genome and environment in the standard gene-based view of inheritance. The focal individual depicted within the gray dashed box has a genome G (dark orange circle) inherited from its parent(s). This genome, when expressed, directs the generation of the phenotype P (dark blue octagon), and the generation of the phenotype is affected by the environment E (green box), which includes both biotic (conspecifics and heterospecifics) and abiotic components. Eventually, the focal individual transmits genomic material G’ to its offspring. What is transmitted, however, is not necessarily identical to what was inherited (G), as a result of mutation and horizontal gene transfer (hence the dark orange arrow from E to G), as well as recombination and meiosis. For the focal individual, an interaction between P and E determines its Darwinian fitness. Solid arrows indicate the role of one component affecting the other, or interactions between components, when an arrow from one component impinges upon an arrow connecting two others. Thus, E interacts with G in generating P, and also affects P directly. Arrows are colour coded to match the colour in which the component they affect is depicted.

In the next section, we briefly review the standard gene-based view of inheritance, emphasizing some points that we will draw upon in order to develop an extended view of inheritance. The subsequent section will be devoted to incorporating non-genetic inheritance into a conceptual framework of how phenotypes are shaped and inherited that will stress the dynamic nature of phenotypes even within the lifetime of an individual. In the final section, we will discuss other studies that have addressed the issue of non-genetic inheritance and its evolutionary implications and identify similarities and differences between our view and theirs, in particular highlighting conceptual distinctions that we believe are important and have not been drawn earlier. We will also discuss some of the broader implications of the extended view of inheritance developed here.

## The standard gene-based view of inheritance

Before developing an extended view of inheritance, we briefly reiterate what the standard gene-based model of inheritance is, and how quantitative genetics distinguishes between the genotypic value and breeding value of a trait in an individual. The fundamental components of this view are that information regarding the specification of phenotypes (whether as blueprint or developmental program) is transmitted to offspring from parent(s) via the inheritance of genomes, and that the specification of the phenotype by the genome can be affected by the environment which includes biotic and abiotic components (Fig. 1). Direct environmental effects on the phenotype of an individual (solid dark blue arrow from E to the octagon around P), however, cannot be passed on to that individual’s offspring, as only genes are transmitted between generations. We note that, in this view, environment in a broad sense can affect the genome (solid dark orange arrow from E to the circle around G in Fig. 1) by inducing mutations or transpositions, or via horizontal gene transfer from conspecifics or heterospecifics. We will not treat acquisition of genetic material by an individual via horizontal gene transfer as constituting inheritance from the viewpoint of that individual, as it does not involve inherited material acquired at conception via transmission from parent(s). Of course, once acquired by horizontal gene transfer, such genetic material can subsequently be transmitted by that individual to its own offspring, becoming part of their inheritance. We next stress a few specific points about this gene-based view of inheritance that we believe will be relevant to the next section entitled ‘An extended view of inheritance’.

i. Although the organism and its phenotype are central in ecology and evolution because Darwinian fitness typically accrues to the individual as a result of its phenotype and how well it functions in a given environmental context, evolutionary dynamics are often tracked at the genetic level. This is because, at least for typical Mendelian traits, the G-P mapping is simple, and the transmission of genes is amenable to mathematical representation. For more complex polygenic phenotypes, a phenotypic approach (quantitative genetics) is taken, although here too there is often an underlying genetic model (see iv, vi below).
ii. The genome plays two distinct roles here: that of a transmissible genome and an expressible genome. During transitions between generations, the genome acts as the transmissible material of heredity, whereas during an individual’s lifetime the genome acts as an expressible material that directs the generation of the phenotype. While there is a degree of materialistic continuity to the genome between generations, functionally the expressed genome can vary temporally and spatially within the lifetime and the body of the individual, respectively (see also v below).
iii. When an individual reproduces, it typically does not transmit the genome it inherited from its parents. This difference between the inherited and transmitted genomes arises due to (a) changes in the genome due to mutations, transpositions and horizontal gene transfer during the individual’s lifetime, and (b) the effects of recombination and meiosis during sexual reproduction.
iv. In the quantitative genetics approach, an individual’s phenotypic value for a trait is conceptualized as being made up of a genotypic value and an environmental effect, with the mean environmental effect assumed to be zero (Falconer and Mackay 1996). The genotypic value is the expected phenotypic value of individuals of a given genotype exposed to all possible environments (Falconer and Mackay 1996). Thus, there is a material partitioning of *causes* of phenotypic value into a genotypic value and an environmental deviation from it. The genotypic value of an individual is, however, not what the individual transmits in terms of phenotypic value propensity to its offspring. Essentially, in a statistical partitioning of *effects*, the genotypic value is further divided into a transmissible component (the breeding value or additive component) and a non-transmissible component traditionally ascribed to genotype-by-genotype interactions. The breeding value of an individual is a measure of how much the mean phenotypic value of that individual’s offspring is expected to differ from the population mean phenotypic value, were that individual to mate at random within the population. In terms of phenotype, the breeding value is what an individual passes on to its offspring, on an average, and it is a function of the individual’s genotype as well as the genotypic composition of the population (Falconer and Mackay 1996).
v. Thus, in terms of the phenotype, the transmissible genome of an individual is reflected in its breeding value (which is dependent also on the genotypic composition of the population) whereas its expressible genome is reflected in its genotypic value, which is independent of the populational context.
vi. The partitioning of phenotypic value into breeding value and a non-transmissible component, and the analogous partitioning of phenotypic variance in a population into a variance of breeding values (the so-called additive genetic variance) and a non-transmissible variance are purely phenotypic, reflecting the degree to which phenotypic differences among individuals are transmitted to the next generation. In this sense, quantitative genetics is essentially a phenotypic theory. Under the assumption of a large number of Mendelian genes contributing to phenotypic value, however, corresponding genetic models can also be developed and have been used in quantitative genetics (Falconer and Mackay 1996).
vii. In this standard gene-based view, the phenotype is conceived of as a largely static entity, at least within the individual’s lifetime.

One last point we wish to make before moving on to developing an extended view of inheritance is that the framework of quantitative genetics, simple though it is, can incorporate many biological phenomena that are often mentioned in the extended evolutionary synthesis literature as being beyond the gene-based view. An early example of such an extended quantitative genetic framework is Cheverud’s (1984) analysis of the evolutionary dynamics of altruistic maternal performance under kin selection in the presence of maternal effects. Similarly, at least from the perspective of *effects* on phenotypic value, an environmental effect and genotype-by-environment interaction reflect phenotypic plasticity and genetic variation for plasticity, respectively. Moreover, many types of adaptive niche construction can be conceptualized as a subset of positive genotype-environment covariance (Saltz and Nuzhdin 2014).

## An extended view of inheritance

We now develop an extended view of inheritance that takes cognizance of phenomena such as transgenerational epigenetic inheritance, parental effects, ecological inheritance and cultural inheritance. We follow the categorization of Danchin et al. (2011); types of non-genetic inheritance have been categorized differently (e.g. Bonduriansky and Day 2009; Helanterä and Uller 2010), but these differences do not affect the development of our view and we defer their discussion to the next section. We first describe some terms and conceptual distinctions we believe to be helpful in discussing such an extended view of inheritance. Our primary focus throughout remains the phenotype: how it develops, what factors affect it, and how it affects other factors. Rather than the distinction between vertical (ancestor to descendent), horizontal (within-generation, between individuals not ancestor-dependent pairs) and oblique (between-generation, between individuals not ancestor-dependent pairs) transmission of information that can affect the phenotype (e,g, Bonduriansky and Day 2009; Helanterä and Uller 2010), we believe it is important to distinguish between heritable phenotypic effects acquired as a material endowment at conception and those acquired subsequently in an individual’s lifetime. Consequently, we will distinguish, in a manner similar to the gene-based view, between inheritome, phenotype and environment while considering how these various factors affect and are affected by the phenotype of a focal individual (Fig. 2). In Fig. 2, time runs from top to bottom and, at conception, the focal individual at the top of Fig. 2 (whose phenotype P_1_ is represented by a dark blue hexagon and inheritome I_1_ by a dark orange circle) inherits its inheritome I_1_ from its parent(s) (dashed dark orange arrow at top of Fig. 2). The inheritome includes DNA sequence, as well as any epigenetic modifications of DNA/chromatin and cytoplasmic mediators of parental effects. The focal individual also receives a phenotype P_1_ as a parental endowment (dashed dark blue arrow at top), comprising, for example, cytoplasmic components and cellular structure. Transiently, the inheritome and phenotype coincide in the fertilized egg. Subsequently, once cell divisions begin, both inheritome and phenotype undergo independent changes and increasingly diverge over time. The environment (outer black box, labelled E in Fig. 2) is conceptualized as a dynamic entity encompassing all individuals and the physical and cultural (if applicable) backdrop within which they live. Clearly, though individuals live and die, the environment has a continuity that, although subject to changes, transcends individual lifetimes: it is a superset to the other categories depicted in Fig. 2. When examining phenotypic variations and their inheritance/transmission from the point of view of a focal individual (Fig. 2), we refer to its local environment (i.e. the subset of E that has a bearing on the phenotype of the focal individual) at a given time step *i* as its contextome at that time step (C*_i_*). Despite their transient conflation in the fertilized egg, we treat the initial phenotype and inheritome as distinct entities because, over the course of a lifetime, the phenotype of an individual undergoes vastly more changes than its inheritome. For example, the phenotype of a multicellular organism changes much more than the inheritome over the course of development from a zygote to an adult. The inheritome, on the other hand, does have considerable integrity across the lifetime. We propose that the terms *inherit* and *transmit* be used solely in the context of an inheritome being received from parent(s) and inheritomic components being passed on to offspring, respectively. We propose using the terms *acquire* and *contribute* to refer to the receiving and effecting of changes to the inheritome and/or phenotype that occur after conception. Of course, changes to the inheritome that are acquired, in this sense, can subsequently be transmitted and therefore inherited by offspring. We further propose a distinction between *parents* and *peers* in terms of this distinction between inheritance and acquisition. Parents transmit phenotypic effects via the inheritome and, transiently, through the initial phenotype, whereas peers contribute phenotypic effects via the inheritome, phenotype and contextome. Biological parents, therefore, can affect offspring phenotypes both as parents, at conception, and as peers, subsequently. This distinction is similar to Bonduriansky and Day’s (2009) use of the term inheritance solely for vertical transmission, and differs from other, broader, usages (e.g. Odling-Smee et al. 2003; Jablonka and Lamb 2005; Helanterä and Uller 2010; Danchin et al. 2011). However, our usage is narrower than that of Bonduriansky and Day (2009) because we restrict inheritance to heritable phenotypic effects acquired as a material endowment at conception.

**Figure 2.**
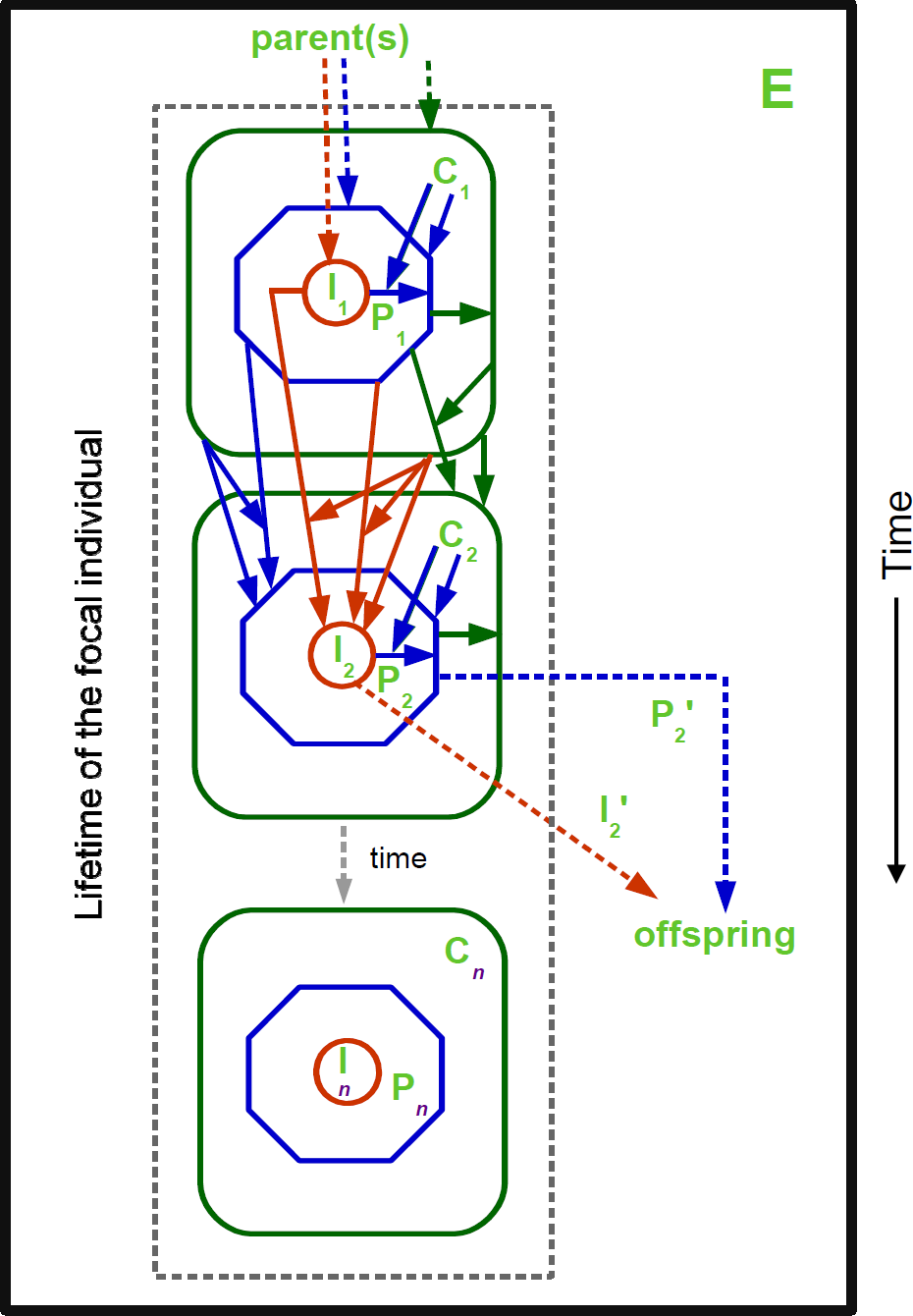
Schematic representation of the relationships between the phenotype P*_i_*, inheritome I*_i_* and environment E, at different time steps (*i* = 1..*n*) during the lifetime of the focal individual depicted within the gray dashed box. The environment (solid dark grey box), which includes all individuals (conspecifics and heterospecifics)as well as the physical and cultural backdrop, is a dynamic superset that transcends the lifespan of individuals of various species. It is the sum of all individuals and together with the physical and, for some species, cultural backdrop within which individuals are transiently embedded during their lifespan. In principle the environment changes on a time-scale that is often similar to the lifespans of individuals. Therefore, the effective local environment of a focal individual is referred to as its contextome (C*_i_* at time step *i*) which changes during an individual’s lifetime. Note that the focal individual is in principle a component of every other individual’s contextome. At conception (time *i* = 1), the focal individual has an inheritome I_1_, inherited from its parent(s) (dashed dark orange arrow). The inheritome includes DNA sequence, as well as any epigenetic modifications of DNA and cytoplasmic mediators of parental effects. The focal individual also receives a phenotype P_1_ as a parental endowment (dashed dark blue arrow); this includes, for example, cytoplasmic components and cellular structure. At conception, the focal individual also inherits and experiences a contextome C_1_ (dashed dark green arrow). In subsequent time steps (2..*n*), the inheritome can change (e.g. from I_1_ to I_2_). This change in inheritome includes changes due to mutation, transposition, horizontal gene transfer and the acquisition of new epigenetic marks due to phenotype, and hence there are solid dark orange arrows from I_1_, P_1_ and C_1_ to I_2_. Solid arrows indicate the role of one component affecting the other, or interactions between components, when an arrow from one component impinges upon an arrow connecting two others. Arrows are colour coded to match the colour in which the component they affect is depicted. Change in the phenotype from P_1_ to P_2_ encompasses changes directed at time step 2 by the expresssed subset(s) of I_2_, subject to interaction with C_2_, and also direct effects of and interactions between P_1_ and C_1_ (dark blue solid arrows). The contextome can also change from C_1_ to C_2_, subject to direct effects of and interactions between P_1_ and C_1_ (green solid arrows). The same set of arrows as between times steps 1 and 2 will apply to every pair of time steps *t* and *t*+1, till *t* = *n*-1, indicated by the dotted gray arrow labeled ‘time’ between time steps 2 and *n*. At any time step *r* (ranging from 2 to *n*), the focal individual may produce offspring, transmitting to them an inheritome I*_r_*’ that is typically different from I*_r_* (dashed dark orange arrows leading to offspring), and a phenotype P*_r_*’, determined by, but different from, P*_r_* (dashed dark blue arrows leading to offspring).

In Fig. 2, the set of arrows between I_1_, I_2_, P_1_, P_2_,C_1_ and C_2_ encompasses the various ways in which inheritome, phenotype and contextome affect phenotypes, and each other, either directly (arrows impinge upon I, P or C) or in an interacting manner (arrows impinge upon other arrows linking I, P or C). How different phenomena sought to be incorporated into an extended evolutionary synthesis correspond to the various direct effects of I_1_, I_2_, P_1_, P_2_, C_1_, and C_2_ on each other is summarized in Table 1. This pattern of potential phenotypic effects of I, P, and C is iterated at every time step, as indicated by the dashed grey arrow labeled ‘time’ connecting I_2_, P_2_, and C_2_ to I*_n_*, P*_n_*, and C*_n_*. Within a given time step, the specific expressed inheritome at that time step directs the generation of the phenotype at that time step, also subject to direct effects of, and interactions with, the contextome at that time step. In contrast to the gene-based view (Fig. 1), however, the phenotype is also directly affected by the phenotype in the previous time step in a manner also subject to interactions with the contextome in the previous time step. Moreover, the phenotype can also be affected directly by the contextome in the previous time step. Thus, in this view, the phenotype during the lifetime of an individual is a dynamic flow from P*_i_* to P*_i_*_+1_, mediated by inputs from the inheritome (I*_i_*_+1_) and the contextome (C*_i_* and C*_i_*_+1_), which themselves are dynamic flows. Thus, the phenotype initiated at conception undergoes transformation throughout the lifetime of the individual under the joint inputs from its previous state and the also changing inheritome and contextome, and interactions between them, in a manner echoing the ‘triple helix’ metaphor of Lewontin (2002).

**Table 1.**
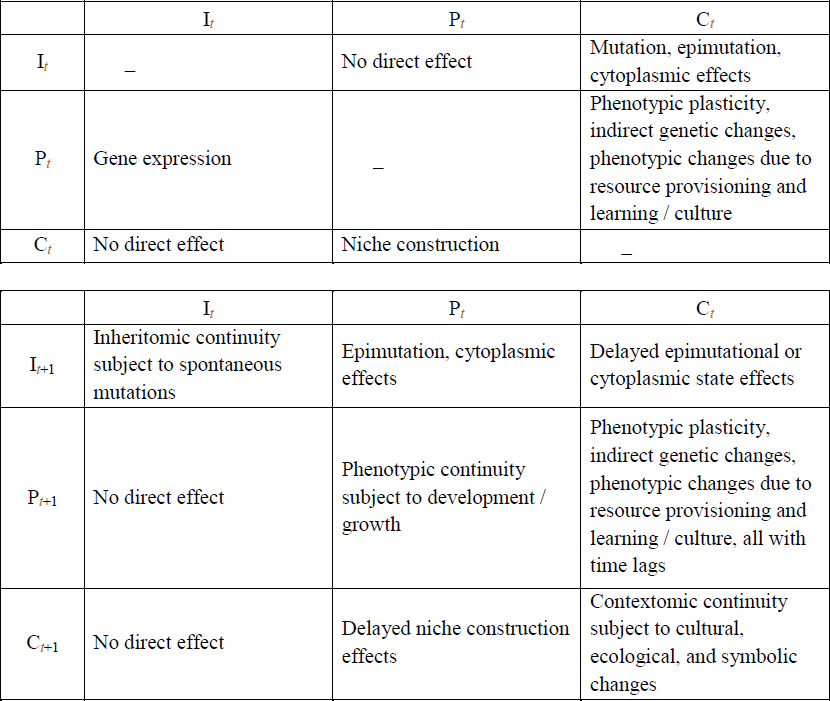
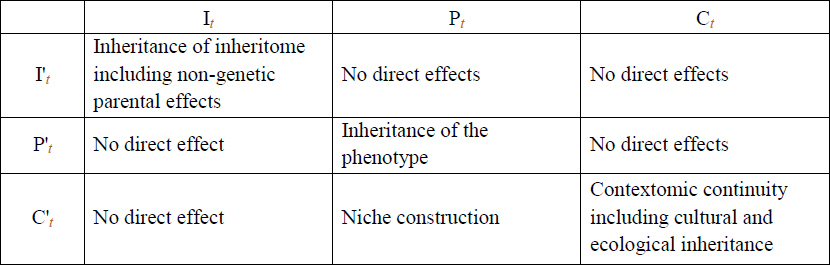
A summary of how the various phenomena sought to be included in an extended evolutionary synthesis are incorporated in the schema outlined in Fig. 2. Entries in the three panels of the table represent direct effects of inheritome (I*_t_*), phenotype (P*_t_*) and contextome (C*_t_*) of a focal individual at time point t on each other (top panel), on the inheritome, phenotype and contextome of the focal individual at the next time point (I*_t_*_+1_, P*_t_*_+1_, C*_t_*_+1_: middle panel), and on the inheritome, phenotype and contextome of an offspring of the focal individual at fertilization for sexually reproducing species and at the point of offspring independence from parental soma for asexually reproducing species (I’*_t_*, P’*_t_*, C’*_t_*: bottom panel). These effects correspond to arrows impinging upon I, P or C in Fig. 2. Indirect effects, for example of C*_t_* on P*_t_*_+1_ via a direct effect of C*_t_* on P*_t_* that, in turn, affects P*_t_*_+1_, are also possible in this schema (arrows impinging upon other arrows linking I, P or C in Fig. 2), but are not depicted in this table.

The inheritome and contextome of the focal individual, in this view, are also conceptualized as flows through time. The inheritome may be altered as a result of its own previous state, and the previous state of the phenotype and contextome, as well as interactions among them (solid dark orange arrows between entities in time steps 1 and 2 in Fig. 2). These changes to the inheritome encompass mutations/transpositions (directed or otherwise), setting or resetting of epigenetic marks on DNA and chromatin, or acquisition of cytoplasmic constituents that may affect offspring phenotypes, mediated by either the physical aspects of the experienced environment, or biological interactions with other individuals. This includes a subset of phenomena included under the label of non-genetic inheritance in the all-encompassing formulations (e.g. Odling-Smee et al. 2003; Jablonka and Lamb 2005; Helanterä and Uller 2010; Danchin et al. 2011). We prefer to treat the remaining mechanisms of non-genetic inheritance, broadly construed, as being changes in the contextome, which is also acquired at conception as a local reflection of the environmental state at that point of time. The contextome of an individual is changing as part of the dynamic change in environment (solid green arrow from C_1_ to C_2_ in Fig. 2). We note that these changes include modifications to the physical or cultural aspects of the environment due to the effects of phenotypes of other individuals, conspecific or heterspecific (e.g. social interactions, competition, niche construction). Thus, contextomic effects on phenotype also include what have earlier been termed indirect genetics effects (Wolf et al. 1998). In addition, an individual’s contextome can be affected by the same individual’s phenotype, subject to interactions with the contextome, and the altered contextome, in turn, can exert effects on subsequent phenotypic states (solid green arrow from P_1_ to C_2_ in Fig. 2). In our view, therefore, we are treating ecological and cultural inheritance (*sensu* Danchin et al. 2011) as being encompassed by the changes to an individual’s contextome and not as inheritance in our narrower usage. Of course, changes made to an individual’s phenotype by its contextome can lead to phenotypic correlations between parent(s) and offspring, as does inheritance, provided the contextome remains the same for the offspring. This is analogous to genotype-environment covariance in quantitative genetics (Salz and Nuzhdin 2014) and we do not believe that it is helpful to label this phenomenon as inheritance, especially since these effects are typically mediated by the phenotypes of many peers that are part of the individual’s contextome, rather than by one or two parents.

During its lifetime, an individual can, in principle, reproduce at any time step. When it does so, it transmits to its offspring, inheritomic material which, in the case of sexual reproduction, together with inheritomic material transmitted by the other parent, form the inheritome of the newly conceived offspring. The parents also pass on an initial phenotype to their offspring which will then undergo changes through the offspring’s lifetime. As was the case for genomes (Fig. 1), what is transmitted/passed on is dependent upon the reproducing individual’s inheritome/phenotype at that time step but is also different from it (distinction between I_2_ and I_2_’, and P_2_ and P_2_’ in Fig. 2). One point of departure from the gene-based view of inheritance here is that inheritomes are more plastic than genomes typically are, since the former are subject to a greater variety of genetic, epigenetic and cytoplasmic changes. Consequently, the source of inheritomic material to be transmitted to offspring by an individual during reproduction at any time step is likely to vary from time step to time step, in contrast to the germline genome which is assumed to be relatively unchanged through life, barring rare mutations.

## Discussion

In this final section, we discuss some implications of the extended view of inheritance developed in the preceding section, especially in the context of earlier attempts to formalize theories of nongenetic inheritance. There are two aspects in which our formulation differs from much of the previous work on this theme. The first is our separation of inheritomic and contextomic mediation of phenotypic correlations between parents and offspring, and our restriction of the term inheritance to describe only the former. The second is our emphasis on the inheritome, and especially the phenotype and contextome, as dynamic flows rather than static entities. We will now discuss some implications of these two emphases.

There are two approaches that have been taken to formalizing the evolutionary consequences of the various types of non-genetic inheritance mechanisms. One is to treat each category – epigenetic inheritance, parental effects, ecological inheritance and cultural inheritance – separately, as is done by Jablonka and Lamb (2005). The other approach is to seek a combined framework within which all forms of non-genetic inheritance can be formalized with regard to their evolutionary implications. This approach has been taken from a quantitative genetics perspective by Bonduriansky and Day (2009) and Danchin et al. (2011), and using the framework of the Price equation by Helanterä and Uller (2010). We note that the important elements of quantitative genetics framework for explaining adaptive evolution can be derived as special cases assuming Mendelian inheritance from the Price equation (Frank 1997). Bonduriansky and Day (2009) treat the change in mean phenotypic value in a population between subsequent generations as being divisible into components due to the change in mean additive genotypic value and in the mean non-genetic component of inheritance. Danchin et al. (2011: Box 4) partition phenotypic variance in the population, V_P_, into transmitted genetic variance (V_G_ in their notation, V_A_ in standard quantitative genetic notation: Falconer and Mackay 1996), transmitted non-genetic variance (V_TNG_) that encompasses phenotypic variance components ascribable to epigenetic, parental non-genetic, ecological and socio-cultural inheritance, and, finally, a non-transmitted component of phenotypic variance. They propose to call the fraction of phenotypic variance that is transmissible [(V_A_ + V_TNG_)/V_P_] the ‘inclusive heritability’ of a trait (Danchin and Wagner 2010) and suggest that it “quantifies the whole evolutionary potential of a trait and can be seen as the result of both genetic and non-genetic heritability” (Danchin et al. 2011). They further suggest that it may be possible to estimate different components of the transmissible phenotypic variance through extensions to classic breeding experiments that also include multiple environmental contexts thought to play a role in non-genetic inheritance, and by tracking epigenetic changes. It is not clear to us that the kinds of partitionings envisaged by Bonduriansky and Day (2009) and Danchin et al. (2011) will actually be experimentally feasible in the context of ecological and cultural inheritance.

Helanterä and Uller (2010) fit various types of non-genetic inheritance mechanisms into the Price equation and try to categorize them into clusters based on the effect each type of non-genetic inheritance has on the different terms in the Price equation (Table 1 in Helanterä and Uller 2010). They find that their approach suggests a clustering different from that into genetic, epigenetic, behavioural and symbolic inheritance (Jablonka and Lamb 2005). They find three clusters that group mechanisms of non-genetic inheritance that will be similarly incorporated into a Price equation framework, and these three clusters are differentiated by whether “inheritance” is by “vertical transmission” (parent to offspring), “induction” (environmentally determined changes between parents and offspring) or “acquisition” (phenotypes affected by peers or other sources). These three clusters differ in the way they mediate adaptive evolutionary change of phenotypes across generations, with and without classic responses to selection in the sense of the breeder’s equation in quantitative genetics (Helanterä and Uller 2010).

Our position is intermediate between the two approaches described above. We note that, of the three clusters of Helanterä and Uller (2010), the “vertical transmission” case corresponds to inheritomic transmission, while their other two clusters are encompassed in contextomic continuity, coupled with contextome effects on phenotype, in our framework. Similarly, the three terms V_G_, V_TEpi_ and V_PNGE_ in the fomulation of Danchin et al. (2011: Box 4), encompassing phenotypic variance due to additive genetic variance, epigenetic variance and parental effect variance, respectively, together correspond to variance due to inheritomic transmission in our framework. What Danchin et al. (2011: Box 4) term “transmitted ecological and socio-cultural variation” is encompassed in contextomic continuity, coupled with contextome effects on phenotype, in our framework. We believe there is a point to separating inheritomic and contextomic effects on the phenotype, for two reasons. First, their effects on adaptive evolutionary dynamics differ, as shown by Helanterä and Uller (2010). Second, following classical quantitative genetics, if we define phenotypic value as being made up of an inheritomic value plus a contextomic value, then it is clear that we need to separately partition each into a transmissible and non-transmissible component. This is exactly analogous to the partitioning of Bonduriansky and Day (2009: Eqns. 2a,b). The partitioning into a transmissible and non-transmissible component of genotypic value is standard in quantitative genetics. Moreover, there is already considerable work on incorporating epigenetic effects, genomic imprinting and parental effects into quantitative genetics models (Kirkpatrick and Lande 1989; Spencer 2002, 2009; Johannes et al. 2008; Tal et al. 2010; Santure and Spencer 2011), suggesting that the partitioning of inheritomic value into transmissible and non-transmissible components is likely to be feasible. It is not clear to us at this time whether a partitioning of contextomic value that is similar in structure to the partitioning for inheritomic value will be possible and, therefore, we believe it might be best to treat it separately. Moreover, we speculate that estimation of the partitioned phenotypic variances due to contextomic and inheritomic effects may require fairly different kinds of experimental designs. A framework analogous to classical quantitative genetics directly suggests possible experimental designs to estimate transmitted versus non-transmitted components of inheritomic variance. For contextomic effects, a completely mechanism free approach like the Price equation may be more suited, with the drawback that it does not lend itself to suggesting designs for experiments.

Our explicit conceptualization of inheritome, contextome and phenotype as flows in time rather than static entities during an individual’s lifetime is another point of departure from many earlier treatments. In particular, this conceptualization suggests that the G-P map metaphor is fundamentally misplaced, as a map is a relation between static entities. In our framework, the inheritome-phenotype map is itself a flow in time and we believe further work is needed to model this relationship more accurately as a dynamic one. One implication of this view of dynamic phenotype is for genome wide association studies (GWAS). It has already been pointed that the problem of “missing heritability” (Maher 2008) is potentially explained if there is a substantial non-genetic component to inheritance in the sense of parent-offspring similarity in phenotype (Danchin et al. 2011). If phenotypes are viewed as changing in time, based largely on previous phenotypic state, modulated by inheritomic and contextomic effects on the phenotype, then it is very likely that, at least for complex traits, the best predictors of phenotypic state maybe previous phenotypic states rather than genotypes. If this speculation is correct, it is a potential explanation for the efficacy of traditional holistic systems of medicine (like Ayurveda in India) in dealing with complex so-called life-style diseases, as these systems try to correlate diseased phenotypic states with a constellation of previous phenotypic states along different trait-axes.

Overall, we believe that the framework for an extended view of inheritance that we have developed here provides a good basis for thinking about specific lines of inquiry relating to the conceptual partitioning of inheritomic and contextomic values, as well as related experimental approaches for obtaining estimates of such partitioned effects and variances. We believe this framework can be fruitfully extended to issues like how non-genetic effects on phenotypes, and their inheritance or acquisition, or phenotypic effects on the contextome (as in niche construction affecting selection pressures: Laland et al. 1999) can affect fitness surfaces; this is something we have not touched upon here because, with the emphasis on inheritome, phenotype and contextome being dynamic flows, the concept of a static fitness surface needs to be critically reexamined. Finally, we believe that the extended view of inheritance and effects on the phenotype developed here is particularly well suited to individual-based simulation studies of evolutionary dynamics incorporating the various possible phenomena discussed in the ongoing extended evolutionary synthesis (Pigliucci and Müller 2010). In particular, given the potential difficulties of designing experiments to deal with partitioning transmitted and non-transmitted components of contextomic effects on phenotypic variance, we believe that data from such individual-based simulations will be very useful for testing the efficacy/limitations of extended quantitative genetic type models, or those based on the Price equation, in terms of how well they can capture the essential features of the evolutionary dynamics of such simulated systems that incorporate all the phenomena used to justify the calls for an extended evolutionary synthesis.

## Acknowledgments

This is contribution no. 1 from the Foundations of Genetics and Evolution Group (FOGEG). FOGEG is an informal association of SD, AJ, NGP and TNCV getting together periodically to work on conceptual issues at the foundations of genetics and evolutionary biology. All authors contribute equally to the manuscripts and the sequence of authors is chosen at random for each submission. The last author acts as corresponding author for that submission. We thank Sachit Daniel and K. P. Mohanan for useful discussions, especially during the early phases of the crystallization of these ideas. We also thank the Indian Academy of Sciences, Bengaluru, for supporting a discussion meeting on ‘Foundations of Evolutionary Theory’ at Orange County, Coorg, in February 2014, at which many discussions on the extended evolutionary synthesis took place. AJ thanks the Department of Science and Technology, Government of India, for support via a J. C. Bose Fellowship. SD, NGP and TNCV thank IISER Pune, IISER Mohali and JNCASR, respectively, for in-house funding.

